# Free-ranging dogs understand human intentions and adjust their behavioral responses accordingly

**DOI:** 10.1101/374595

**Authors:** Debottam Bhattacharjee, Shubhra Sau, Anindita Bhadra

## Abstract

Domestic dogs (*Canis lupus familiaris*) are remarkably sensitive and responsive while interacting with humans. Pet dogs are known to have social skills and abilities to display situation-specific responses, but there is lack of information regarding free-ranging dogs which constitute majority of the world’s dog population. Free-ranging dogs found in most of the developing countries interact constantly with familiar and unfamiliar humans receiving both positive and negative behavior. Thus, understanding human intentions and subsequent behavioral adjustments are crucial for dogs that share habitats with humans. Here we subjected free-ranging dogs to different human social communicative cues (friendly and threatening – low and high), followed by a food provisioning phase and tested their responsiveness. Dogs exhibited higher proximity seeking behavior as a reaction to friendly gesture whereas, they were prompted to maintain distance depending on the impact of the threatening cues. Interestingly, only the high-impact threatening showed to have a persistent effect which also remained during the subsequent food provisioning phase. An elevated approach in the food provisioning phase elicited the dependency of free-ranging dogs on humans for sustenance. Our findings suggest that free-ranging dogs demonstrate behavioral plasticity on interacting with humans; which provides significant insights into the establishment of the dog-human relationship on streets.

## INTRODUCTION

Recent trends in research on interspecific interactions have unveiled several important aspects regarding the interplay of the component species. Investigating the eco-ethology of one component species and its trajectories can provide adequate information on the other (Bertness and Callaway, 1994; Thompson, 1999). Human-animal interaction is one such field that attracts researchers to find solutions for evolving problems like human-animal conflict, spread of zoonoses, uncontrolled population growth of unwanted species, etc. In the recent times, studies on human-animal interactions have enabled us to apprehend evolutionary processes like domestication (Hare et al., 2002; Miklósi and Soproni, 2006). Such scientific investigations, coupled with comparative analyses, also helped us understand the functionality of behaviors and communicative intents of species. As the first domesticated species, dogs have spent a considerably long period of time socially interacting with humans (Larson et al., 2012; Morey, 2006; Perri, 2016). Thus, exploring the dog-human interaction paradigm, is specifically helpful to analyse the underlying dynamics of the domestication process that enabled the transition of wolf-like ancestors to man’s best friend.

Domesticated dogs interact with humans regularly and possess social abilities to respond to various human actions (Hare and Tomasello, 2005; Miklósi and Soproni, 2006). Dogs are highly sensitive to human communicative cues like pointing, touching, body orientation etc. and in utilising such cues to find hidden rewards such as food (Hare et al., 2002; Miklósi and Soproni, 2006). It has been suggested that domestication played a pivotal role in the development of human-like social skills in dogs (Hare and Tomasello, 2005). At the same time, substantial evidence for the importance of life history and ontogenic experience with humans in the development of the dog-human relationship is also present (Dorey et al., 2010; Wynne et al., 2008). Dogs have been shown to flexibly adjust their behavior in several interactive instances with humans such as, avoiding pointing cues provided by “unreliable humans” (Takaoka et al., 2015), following pointing cues only on being rewarded in a preceding trial, thereby showing the ability to adjust behavioral responses (Bhattacharjee et al., 2017a) etc. One study also confirmed dogs’ understanding of human attentional states, suggesting the fact that dogs can specifically ask for help when a human is paying attention to it (Miklósi et al., 2000). Dogs have been shown to give emphasis on human body and face while decoding intentions (Nagasawa et al., 2011). Vas et al. (2005) found that pet dogs can differentiate between friendly and threatening cues provided by an unfamiliar human and can display situation-relevant behavior. The same study also reported breed specific differences of dogs’ responsiveness due to varying levels of sensitivity towards humans. Except for one study mentioned above, all the other studies explored behavioral plasticity in pet dogs and hence is not representative of dogs that are not under the direct supervision, and thereby influence, of humans.

Free-ranging dogs that are primarily present in the developing countries make up almost 80% of the world’s dog population (Boitani and Ciucci, 1995; Hughes and Macdonald, 2013). However, studies are significantly lacking and insufficient with these dogs. They are partially dependent on humans for their sustenance, but their activities are not directly controlled by humans (Bhadra et al., 2016; Sen Majumder et al., 2014; Paul et al., 2016). Unlike pet dogs, the free-ranging subpopulation receive both positive and negative human influences; humans play the most significant role in mortality of these dogs (Paul et al., 2016), while also being the primary provider of food. This particular difference, along with other ecological parameters pertaining to survival (competition for food, inter-group dynamics etc.) make free-ranging dogs different from pets. The interaction between free-ranging dogs and humans on the streets are quite complex and dynamic. They usually avoid human contact but can form strong bonds over repeated positive interactions with unfamiliar humans (Bhattacharjee et al., 2017b). Additionally, they need to understand and decipher human intentions clearly. A recent study showed that very young pups of free-ranging dogs follow simple human pointing cues but learn to adjust the behavior when they grow up and start foraging events. A greater risk of negative impact from humans like beating, harassment and threatening are probably the prime reasons for such plasticity in point-following behavior (Bhattacharjee et al., 2017a). However, this study does not provide us insights into the dogs’ understanding of social cues that are used by humans in day to day interactions with these dogs.

Here we used three different types of commonly used human social communicative cues while interacting with dogs on streets. The cues differed in terms of their actions and representations. The friendly cue illustrated an affiliative gesture, while the low and high impact threatening cues had negative display of gestures. In addition to the three cues, we used a neutral cue as control. We tested the responses of the dogs to the cues and investigated the influences in a post-cue food provision phase. We hypothesize that the dogs would enact positively upon receiving the friendly cue showing higher proximity and approach behavior. Additionally, the dogs would avoid human proximity and adjust their responses flexibly according to the impacts of the two threatening cues.

## MATERIALS AND METHODS

### Subjects and study area

We tested 120 adult, physically fit, solitary free-ranging dogs. Dogs were located randomly in different areas of West Bengal, India (see Supplementary **Figure S1**) for the experiment - Mohanpur (22°56’49’’N and 88°32’4’’E), Kalyani (22°58’30”N, 88°26’04”E), Sodepur (22°69’82’’N and 88°38’95’’E) and Kolkata (22°57’26”N, 88°36’39”E). Sexes of the dogs were determined by observing their genitals. To rule out any possibility of resampling, we tested dogs from different locations on different days and photographed each individual to collect information on coat color, scar marks and other morphological features.

### Experimental Procedure

We used four different experimental conditions pertaining to various social cues in order to investigate the response of solitary free-ranging dogs towards an unfamiliar human. We tested separate sets of 30 dogs in each of the four experimental conditions. All the experimental trials were conducted on the same locations where the focal free-ranging dog was found (e.g. streets, markets, residential areas etc.). One piece of raw chicken (10-12g) was used as food. The experimenters (E1 and E2) were consistent throughout the study, played specific roles and were young males. Video recording was done from a distance using a Sony HDR-PJ410 camera mounted on a tripod.

The experimental conditions comprised of 2 major and 3 minor phases, carried out in the following order (Figure **S2**) –

#### Attention seeking phase (minor)

E2 attracted the attention of a solitary dog by making very short vocalization for 1-2 seconds (Bhattacharjee et al., 2017a). This step was necessary as we found some dogs lying down, resting or dozing. To keep the protocol consistent, E2 carried out this step in all the four experimental conditions.

#### Transition phase (minor)

Once the dog was alerted, E2 immediately left the place and stood behind the camera, which was kept at a minimum distance of 4.5 m from the dog. E1 arrived near the position where E2 was initially standing. This whole procedure was completed within 10 seconds.

#### Social cue phase / SCP (major)

E1 stood approximately 1.5 m away from the dog, facing it. Since the dogs were free-ranging and not on leash, E1 had to adjust his position in order to maintain the approximate distance of 1.5 m. After standing at the specified spot, E1 provided any of the predetermined social cue for 30 seconds –

- *Friendly Cue (FC) -* E1 displayed a positive gesture by bending slightly forward and extending both the arms (see Supplementary **Movie S1**). In India, people use similar gestures (sometimes clubbed with positive vocalisations) to provide positive social rewards to dogs. E1, while providing the social cue, gazed and tried to maintain eye-contact with the dog. E1 did not touch the focal dog deliberately in order to avoid any potential bias of social contact.
- *Low impact threatening (LIT) -* E1 raised one of his hands (counterbalanced) and gazed at the dogs (see Supplementary **Movie S2**). The cue differed from FC in having a negative display of human gesture. People on the streets often raise one of their hands to scare, threaten or shoo away dogs. We have adopted the same gesture in our protocol to investigate the effects and associated responses.
- *High impact threatening (HIT) -* This phase differed from LIT in terms of impact. Here E1 used a 0.45 m long solid wooden stick in his hand (counterbalanced) while providing the gesture (see Supplementary **Movie S3**). E1 had to hide the wooden stick in the transition phase before enacting the gesture.
- *Neutral Cue (NC) -* Here, E1 stood in a neutral posture and looked straight ahead and did not enact any gesture.

#### Food transfer phase (minor)

Immediately after SCP, food was provisioned. E2 came quickly, handed over the food to E1 and went back to his position behind the camera. The process was completed within 10 seconds and care was taken to ensure that the focal dog did not see the transfer of food to E1.

#### Food provisioning phase / FPP (major)

E1 again adjusted his position to keep the distance consistent and placed the food on the ground. The food was placed at a distance of 0.3 m from E1, thus at a distance of 1.2 m from the dog. E1 stood in a neutral position after placing the food and looked straight ahead, without making eye contact with the dog (see Supplementary **Movie S4**). FPP lasted 30 seconds or until the dog obtained the food, whichever was earlier. Food was removed in case a dog did not obtain it.

Except for the SCP, all the other phases were constant and exactly similar across the experimental conditions.

### Data Analysis and statistics

We coded all the important behaviors relating to the experiment, which have been listed in the ethogram below (**Table 1**).

**Table 1.** List of behaviors coded from the videos and their definitions.

| Phase | Behavior |  | Definition |
| --- | --- | --- | --- |
| SCP | Approach | | Subject moved towards E1, distance between E1 and subject was $\leq 0.3$ m. |
|  | No approach | Same position | Distance between E1 and subject was equal to 1.5 m. |
| | | Distant | Distance between E1 and subject was $> 1.5$ m. |
|  | First reaction |  | The first behaviour observed as a reaction to the social cue – gazing, gazing with tail wag, scared and moving back, no reaction. |
|  | Demeanor (Holistic) | Affiliative | All behaviors towards E1 that are involved in the formation and maintenance of human-dog bonding. Includes attention seeking, proximity seeking, contact seeking, social facilitation, tail wagging, and relaxed posture (Rehn et al., 2013). |
|  |  | Aggressive | Agonistic and/or aggressive (dominant or threatening) behavior towards E1. Includes mouthing, biting clothing, jumping up, snapping, growling, baring the teeth, stiff posture, staring and/or "whale-eyeing," high, stiff tail carriage, piloerection (Overall, 2014). |
|  |  | Anxious | Anxious or fearful behavior towards E1. Includes shaking (trembling), excessive panting, lip-licking, urination, tail between the legs, running away, flinching, corners of the mouth retracted down and back. May be maintaining distance from the E1 (Lindsay, 2005). |
|  |  | Neutral | Demeanor that is not otherwise covered in this ethogram. May include resting and sleeping during the experiment, exploratory behavior not directed at E1 or food (sniffing, digging, chewing, scent rolling), self-care (scratching, licking), or general disinterest in E1. |
| | Human Proximity | | Distance between E1 and subject was $\leq 0.3$ m. |
|  | Gazing at human |  | Subject is sitting, standing, or lying and focused on (muzzle turned towards) E1's body or face. Cumulative duration of gazing / looking behavior at E1. |
| FPP | Approach | | Subject moved towards E1, distance between food and subject was $\leq 0.3$ m. |
|  | No approach | Same position | Distance between food and subject was equal to 1.2 m. |
| | | Distant | Distance between food and subject was $> 1.2$ m. |
|  | Latency |  | Time taken to obtain the food after its provision on the ground. Valid only for subjects that obtained food. |
| | Feeding time (proximity) | | Time taken to eat the food in front of E1. Distance of subject and human should be $\leq 0.3$ m. Distant ($> 0.3$ m) feeding or taking food away was not considered. |
|  | Gazing at human |  | Same as social cue phase. |

## RESULTS

Various statistical tests were carried out for the analysis. Since we have compared parameters of the major phases in all possible combinations, description of some of the post-hoc statistical tests were presented in supplementary material to avoid congestion in the main text.

### Number approached

Dogs approached differently in SCP and FPP of the four conditions (Contingency χ^2^: χ^2^ = 10.439, df = 3, p = 0.015, **Figure 1**). In the NC condition, initially 4 individuals approached, while the number increased to 17 in FPP, the change being statistically significant (Goodness of fit χ^2^: χ^2^ = 8.048, df = 1, p = 0.005), thereby indicating a distinct positive impact of food. However, we did not find any difference between the two phases in the FC condition as dogs equally responded to both positive gestures (25) and food (30), more than expected by chance alone (Goodness of fit χ^2^: χ^2^ = 0.455, df = 1, p = 0.50). The LIT condition had a very momentary impact as only 1 individual approached in SCP, while 13 individuals approached in FPP (Goodness of fit χ^2^: χ^2^ = 10.246, df = 1, p = 0.001). Thus, dogs flexibly adjusted their behavior and tended to approach more when food was provided. Unlike the LIT condition, we found a strong effect of HIT, where none of the individuals approached initially and only 1 (Goodness of fit χ^2^: χ^2^ = 1.000, df = 1, p = 0.317) in the later phase, when the food reward was offered. This was suggestive of the dogs’ perception of human intentions based on an immediate encounter.

**Figure 1.**
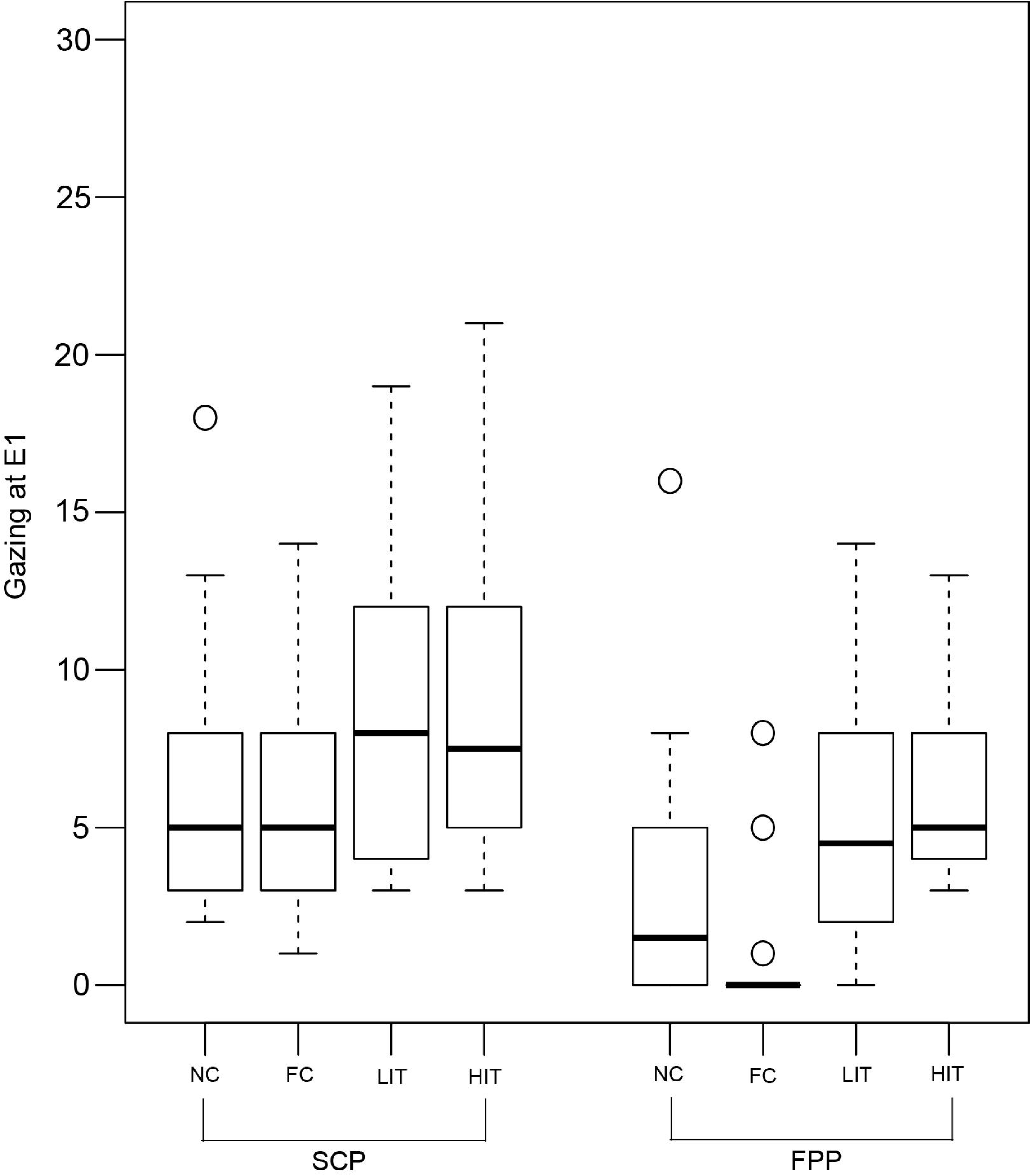
Number of approaches. Bar graph showing the number of approaches in the SCP and FPP of the four experimental conditions – NC, FC, LIT and HIT. Number of dogs that approached varied between the phases across the conditions (Contingency χ^2^: χ^2^ = 10.439, df = 3, p = 0.015). Asterisks indicated significant differences. The dotted line indicated the chance level (50%).

We compared the number of approaches across conditions for both SCP and FPP. Significantly higher number of dogs approached in SCP in response to FC, as compared to the NC, LIT and HIT conditions (see Supplementary **Table S1**). We noticed a marginal difference between the NC and HIT conditions (see Supplementary **Table S1**). Additionally, the number of approaches in the SCP of LIT did not differ from NC and HIT conditions (see Supplementary **Table S1**).

Comparison of the number of approaches among FPP of the four conditions revealed interesting results. Dogs approached significantly more in FC compared to LIT (Goodness of fit χ^2^: χ^2^ = 6.721, df = 1, p = 0.010) and HIT (Goodness of fit χ^2^: χ^2^ = 27.129, df = 1, p < 0.0001), but not NC (Goodness of fit χ^2^: χ^2^ = 3.596, df = 1, p = 0.058), again implying the role of the food provisioned. In the HIT condition, the number of dogs that approached was significantly lower than the LIT condition (Goodness of fit χ^2^: χ^2^ = 10.286, df = 1, p = 0.001), suggesting an influence of the HIT cue which even surpassed the impact of food. In addition, we also found that the HIT cue differed from NC, but the number of approaches in FPP of LIT and NC conditions were comparable (see Supplementary **Table S1**). These results together reinforce the idea that dogs were capable of differentiating between the high and low impact threatening cues, and act accordingly, to maximize their chances of obtaining the food reward while avoiding serious threat.

### No approach

We calculated the number of individuals in different conditions that did not approach and further divided the numbers in two subcategories – ‘same’ and ‘distant’ position (see **Table 1**). Here we laid emphasis on the ‘distant’ position which served as a correlate of negative impact. Consistent with our hypothesis, we could not see any dog running or moving away in the NC and FC conditions, thereby dogs exclusively showed no approach of ‘same position’ subcategory. Thus, we analysed the data only from LIT and HIT conditions. We used the percentage of responses out of the total “no approach” cases for all the comparisons.

52% and 24% of the dogs were distant in SCP and FPP respectively, in the LIT condition (Goodness of fit χ^2^: χ^2^ = 10.316, df = 1, p = 0.001, **Figure 2**). Consistent with this, we also found dogs showing significantly more distant positions in SCP (73%) than the FPP (45%) of the HIT condition (Goodness of fit χ^2^: χ^2^ = 6.644, df = 1, p = 0.01, **Figure 2**). Further comparisons revealed a significant difference between the FPP of LIT and HIT conditions (Goodness of fit χ^2^: χ^2^ = 6.391, df = 1, p = 0.011, **Figure 2**), where higher numbers of dogs stayed at the ‘distant’ position in the HIT condition. However, we did not find any difference between the SCP of the two conditions (Goodness of fit χ^2^: χ^2^ = 3.528, df = 1, p = 0.060).

**Figure 2.**
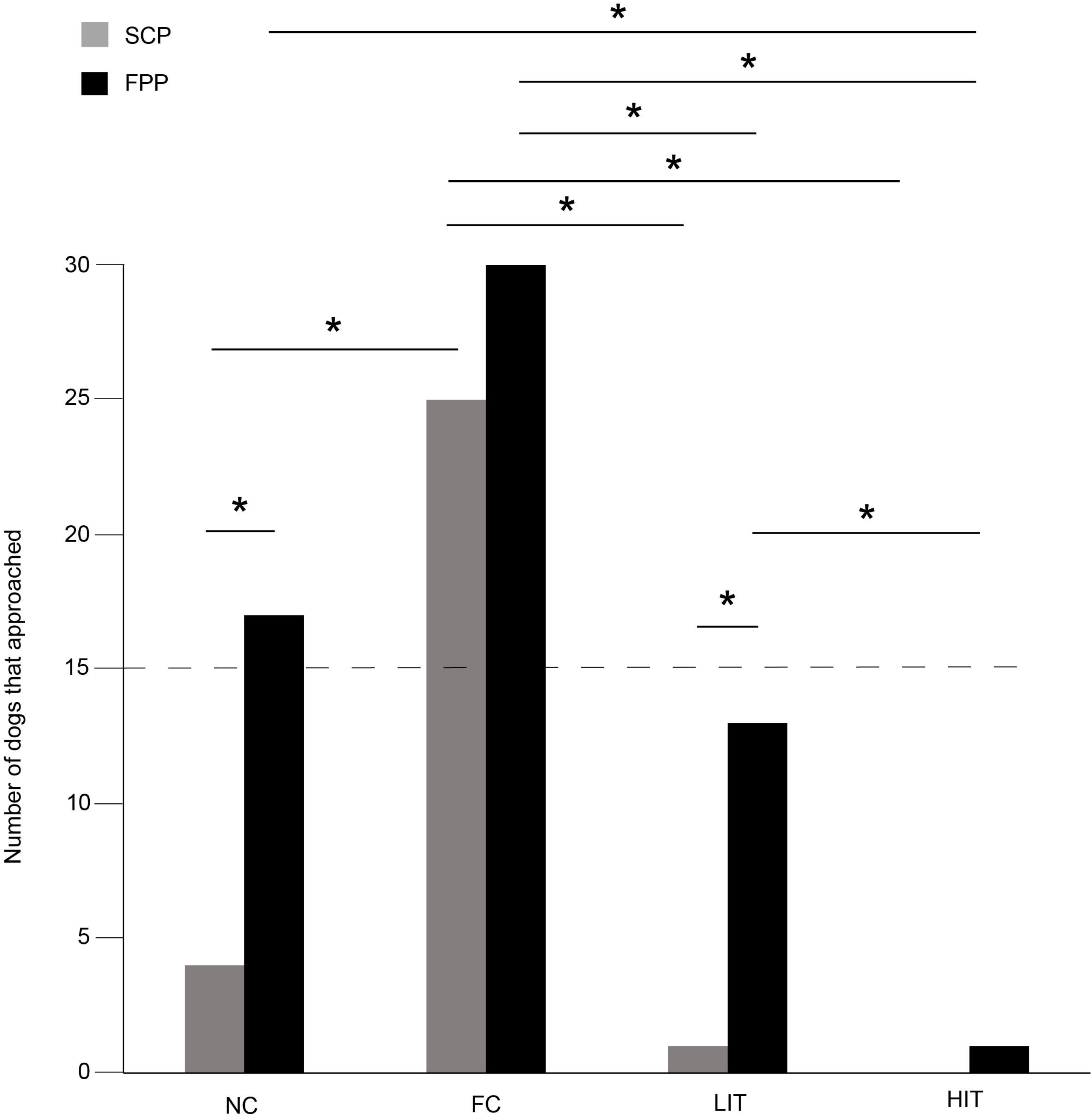
Percentage of distant position out of “no approach”. Bar graph showing the percentage of dogs that showed distant position out of “no approach” in the SCP and FPP of LIT and HIT conditions. Dogs showed significantly more distant positions in both the SCP compared to FPP of LIT (Goodness of fit χ^2^: χ^2^ = 10.316, df = 1, p = 0.001) and HIT (Goodness of fit χ2: χ2 = 6.644, df = 1, p = 0.01) conditions. Asterisks indicated significant differences.

### First reaction to a social cue

Quantification of the first reaction was important in terms of impact and effect of the social cues. We found that the reactions (see **Table 1**) were distributed differently in the four experimental conditions. In the NC condition, dogs showed varying levels of reactions. 60% of the dogs showed gazing behavior, 10% showed gazing with tail wagging, 30% stayed neutral and displayed no particular reaction. None of the dogs showed a fear response. We found a significant difference among the proportion of individuals showing the different reactions (Goodness of fit χ^2^: χ^2^ = 25.200, df = 3, p < 0.0001, **Figure 3A**). Gazing and no reaction were comparable and displayed at a higher rate than other behaviors (see Supplementary **Table S1**). In the FC condition, we found 80% of the dogs showing gazing with tail wagging as their first reaction, while 20% showed gazing behavior only. No dog showed a fear response, and all dogs responded. Gazing with tail wagging occurred at a significantly higher rate than only gazing behavior (Goodness of fit χ^2^: χ^2^ = 10.800, df = 1, p = 0.001, **Figure 3B**). Except for a single individual, all the dogs reacted in the LIT condition. 60% of the dogs showed fear response to the social cue at a significantly higher rate than both gazing (20%) and gazing with tail wagging (17%) behaviors (Gazing – Goodness of fit χ^2^: χ^2^ = 6.000, df = 1, p = 0.014; Gazing with tail wagging – Goodness of fit χ^2^: χ^2^ = 7.348, df = 1, p = 0.007, **Figure 3C**). In the HIT condition, 97% of the dogs showed fear response when the threatening gesture was enacted, whereas only one individual displayed gazing with tail wagging (Goodness of fit χ^2^: χ^2^ = 26.133, df = 1, p < 0.0001, **Figure 3D**).

**Figure 3.**
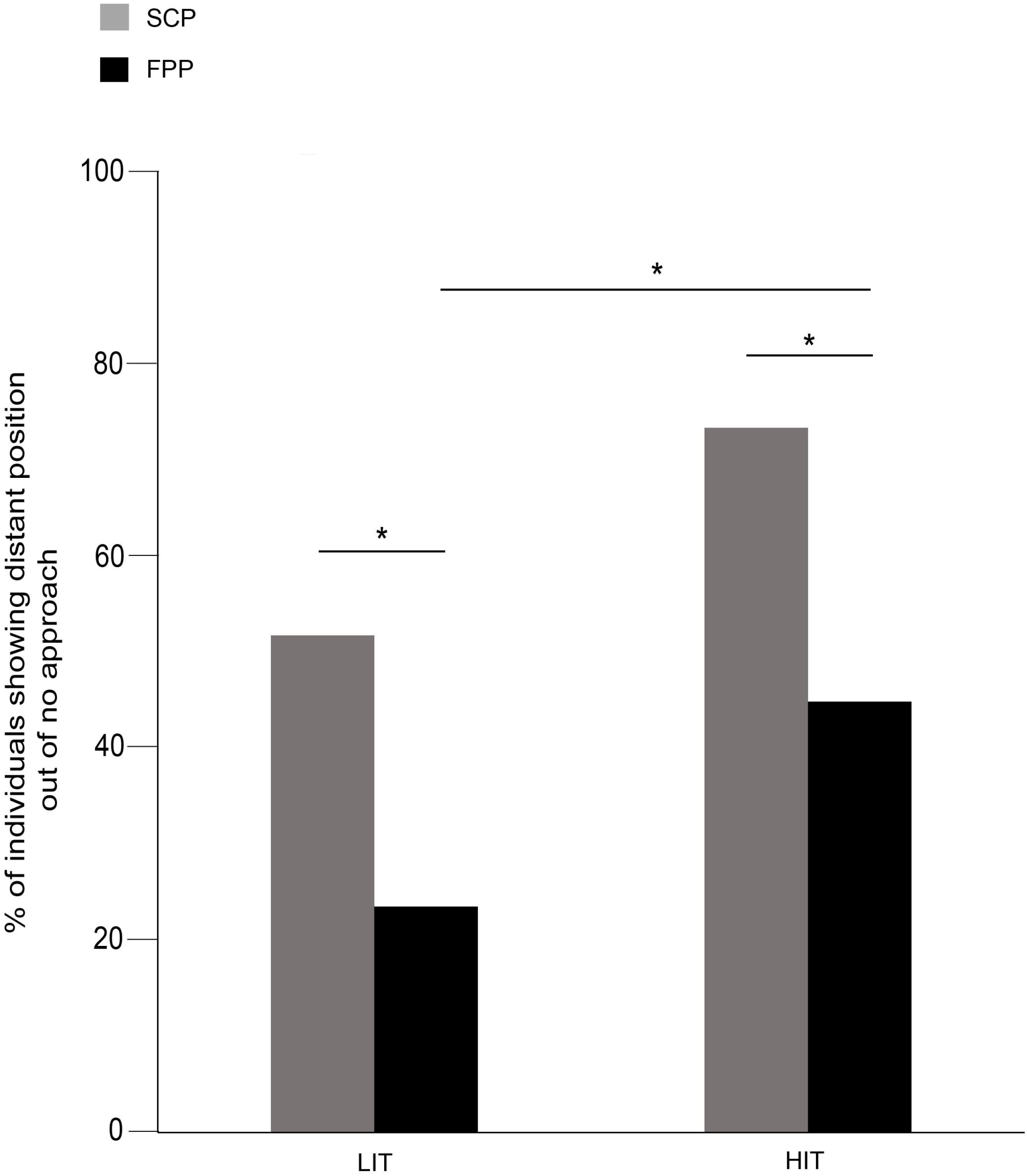
First reaction to social cue. Pie charts showing the first reaction (behaviors) to social cues. (A) Distribution of behavioral responses in NC condition, (B) Distribution of behavioral responses in FC condition, (C) Distribution of behavioral responses in LIT condition, (D) Distribution of behavioral responses in HIT condition.

### Demeanor

Dogs displayed mostly neutral (43%) and anxious (43%) behaviors in the NC condition. Affiliative behaviors were shown at a lower rate than both the neutral and anxious behaviors (Goodness of fit χ^2^: χ^2^ = 4.765, df = 1, p = 0.029, **Figure 4**). Agonistic or aggressive behaviors were absent. Unlike the outcomes in NC, majority of dogs (80%) showed affiliative behaviors, rather than neutral (Goodness of fit χ^2^: χ^2^ = 12.448, df = 1, p < 0.0001) and anxious behaviors (Goodness of fit χ^2^: χ^2^ = 21.160, df = 1, p < 0.0001) in FC. Aggression was not observed. In the LIT condition, 57% of the dogs showed anxious behaviors, which was higher than all the other three categories - (Neutral – Goodness of fit χ^2^: χ^2^ =6.545, df = 1, p = 0.011; Affiliative - Goodness of fit χ^2^: χ^2^ = 6.545, df = 1, p = 0.011; Aggressive – Goodness of fit χ^2^: χ^2^ = 9.800, df = 1, p = 0.002). 97% of the dogs showed anxious behaviors in HIT condition. However, we did not see a statistical difference between the levels of anxious behavior shown in LIT and HIT conditions (Goodness of fit χ^2^: χ^2^ = 3.130, df = 1, p < 0.07).

**Figure 4.**
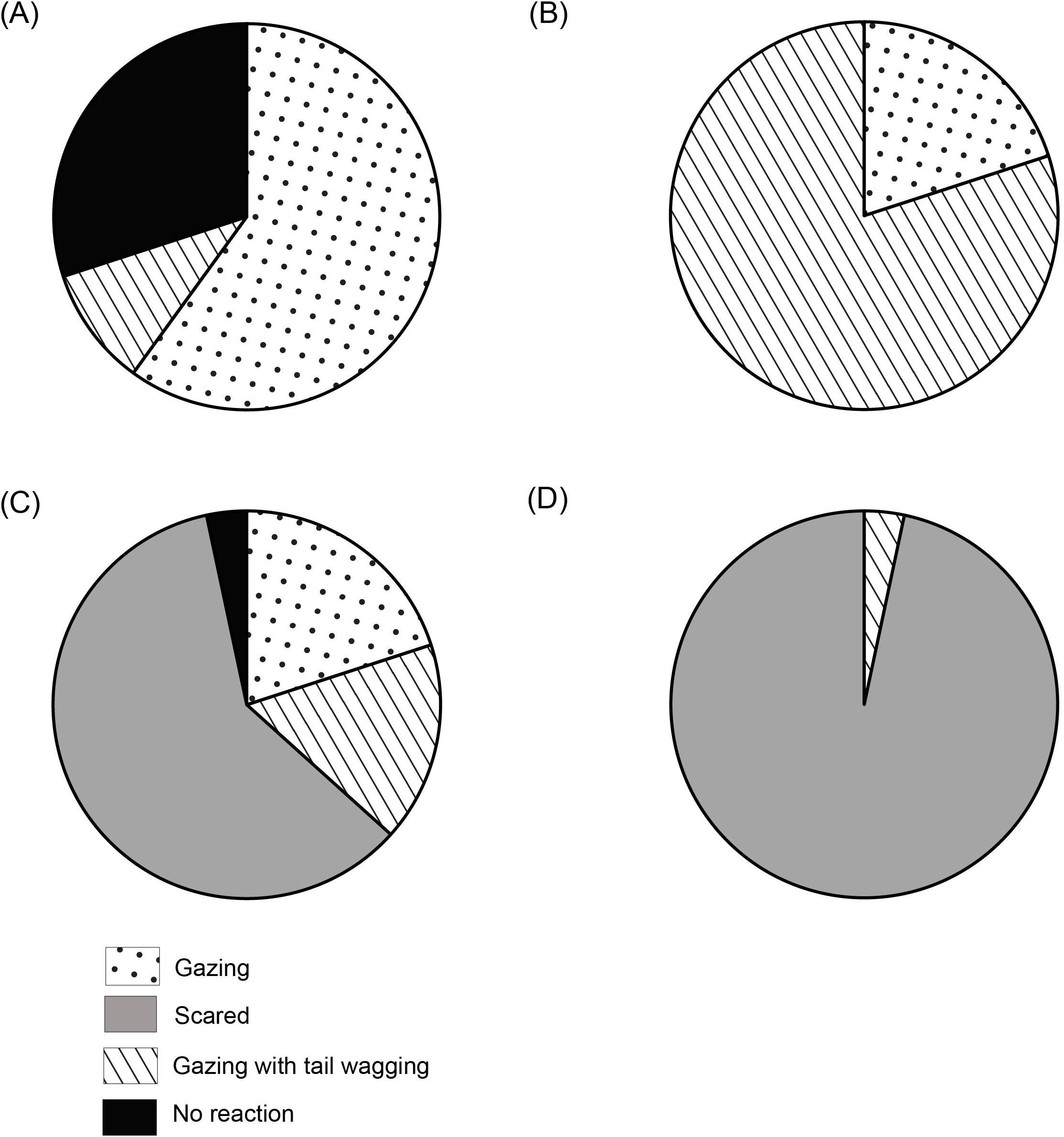
Demeanours (Holistic). Stacked bar graph showing the demeanours of dogs in SCP of NC, FC, LIT and HIT conditions.

### Human proximity

Dogs showed varying levels of proximity to the human experimenter in the different conditions (Kruskal – Wallis test, χ^2^ =77.127, df = 3, p < 0.0001, **Figure 5**). Post-hoc pairwise comparisons revealed that the duration of human proximity was higher in the FC condition compared to others (see Supplementary **Table S1**). However, we did not find any difference between the duration of human proximity in the NC, LIT and HIT conditions (see Supplementary **Table S1**), indicating a general avoidance of human proximity in free-ranging dogs.

**Figure 5.**
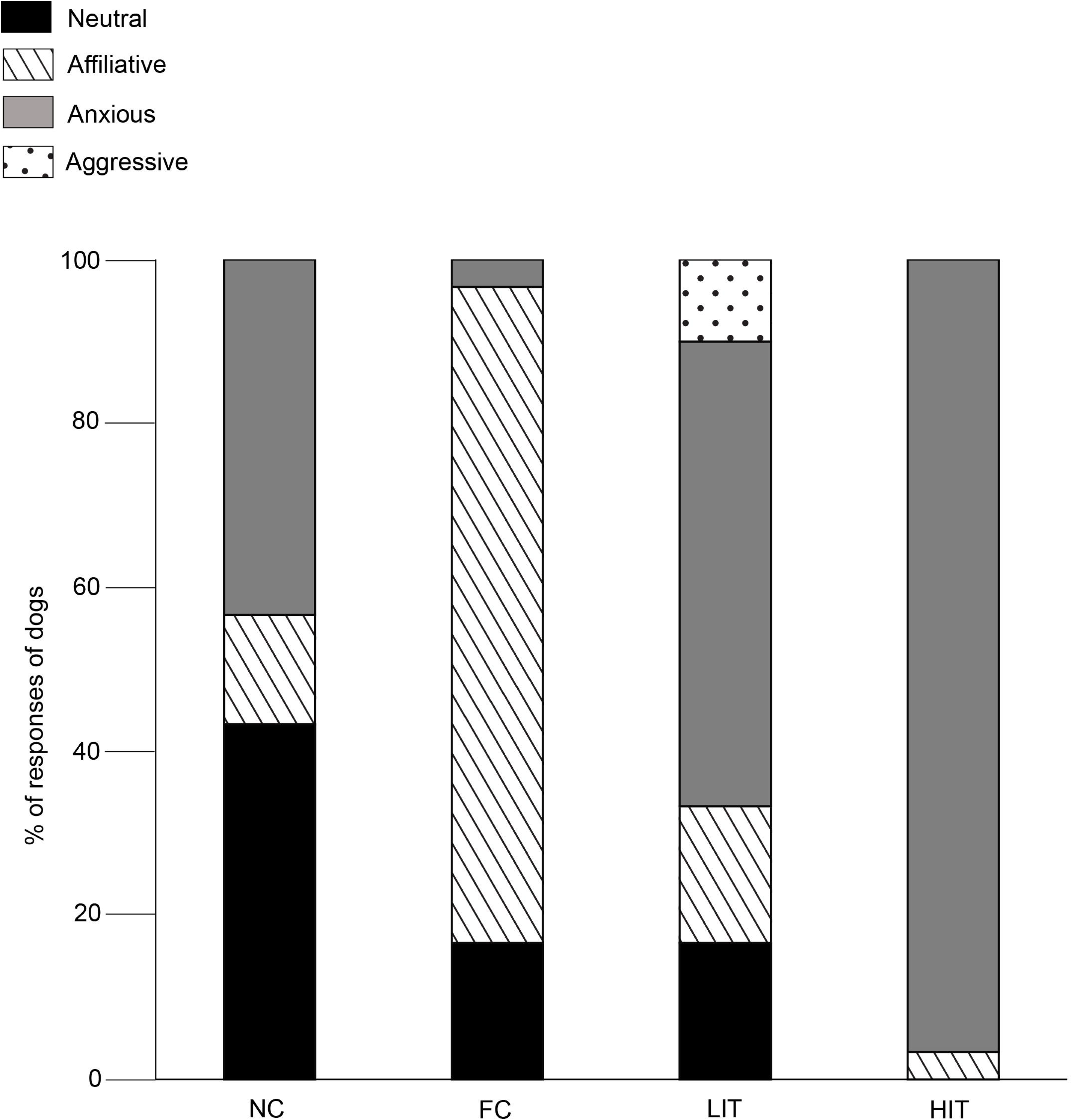
Duration of human proximity. Box and whisker plot illustrating the duration of human proximity of dogs in SCP of NC, FC, LIT and HIT condition. Dog showed significantly higher proximity to E1 in the FC (12.43±9.54 sec) than in other conditions. Boxes represent interquartile range, horizontal bars within boxes indicate median values, and whiskers represent the upper range of the data. “a” and “b” indicate significant differences.

### Gazing

GLM analysis revealed that both the LIT and the HIT conditions are significant predictors of gazing at E1 in the SCP (**Figure 6**, **Table 2**).

**Figure 6.**
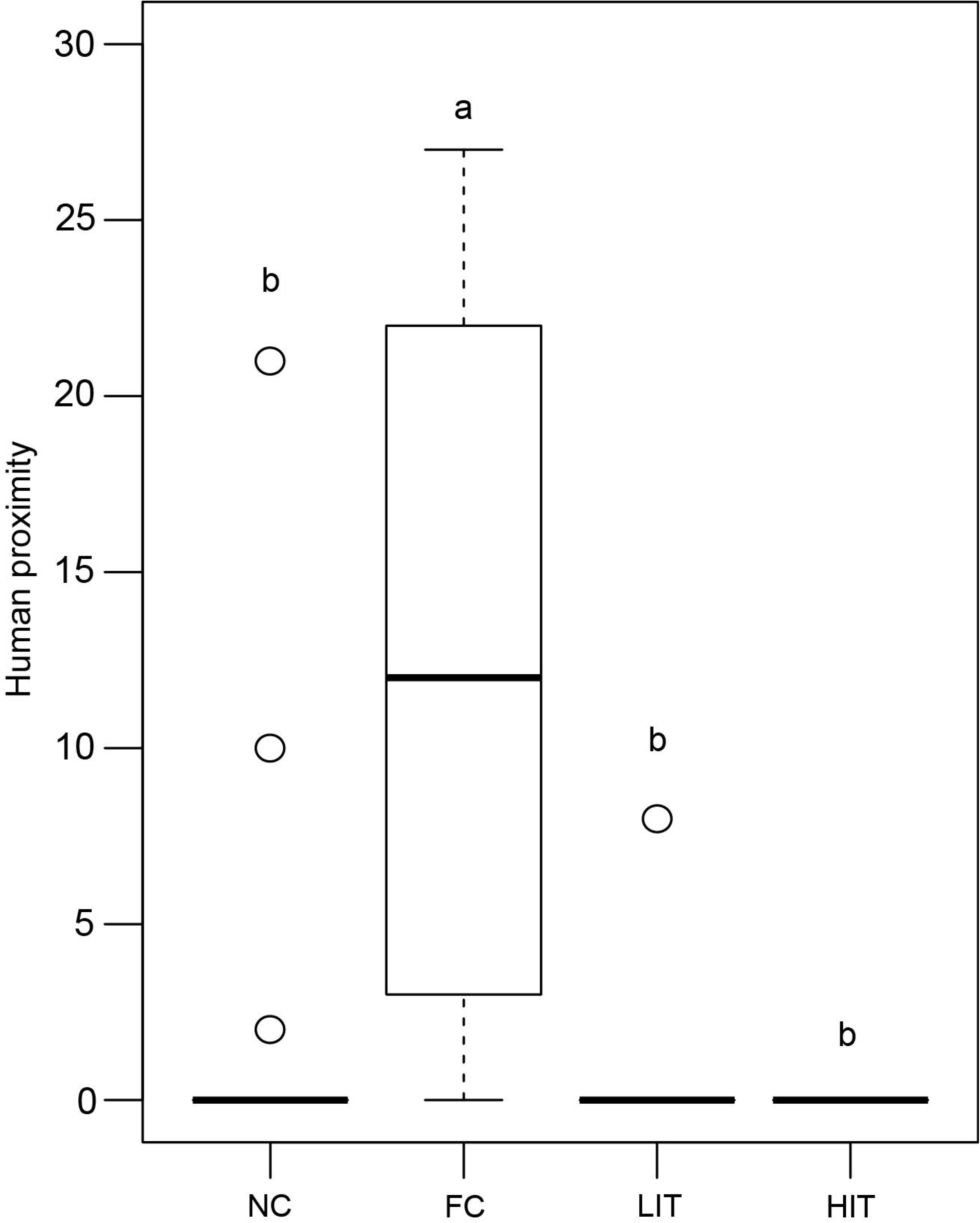
Duration of gazing. Box and whiskers plot showing the duration of gazing at E1 in SCP and FPP of NC, FC, LIT and HIT conditions. Boxes represent interquartile range, horizontal bars within boxes indicate median values, and whiskers represent the upper range of the data.

**Table 2.** GLM results showing the effect of experimental conditions on gazing behavior in the SCP.

|  | estimate | Standard error | z- value | Pr(>\|z\|) |
| --- | --- | --- | --- | --- |
| fixed effects |  |  |  |  |
| Intercept | 1.75786 | 0.07581 | 23.188 | < 2e-16 *** |
| Condition FC | 0.01143 | 0.10691 | 0.107 | 0.914865 |
| Condition HIT | 0.43937 | 0.09722 | 4.520 | 6.2e-06 *** |
| Condition LIT | 0.37828 | 0.09841 | 3.844 | 0.000121 *** |

Interestingly, in the FPP, we found all the different conditions to be significantly contributing to the prediction of the duration of gazing behavior (**Figure 6**, **Table 3**). Dogs gazed the least (0.46 ± 1.69 sec) in the FPP of the FC condition.

**Table 3.** GLM results showing the effect of experimental conditions on gazing behavior in the FPP.

|  | estimate | Standard error | z- value | Pr(>\|z\|) |
| --- | --- | --- | --- | --- |
| fixed effects |  |  |  |  |
| Intercept | 1.1206 | 0.1043 | 10.748 | < 2e-16 *** |
| Condition FC | -1.8827 | 0.2869 | -6.563 | 5.28e-11 *** |
| Condition HIT | 0.6315 | 0.1290 | 4.894 | 9.88e-07 *** |
| Condition LIT | 0.4822 | 0.1326 | 3.636 | 0.000277 *** |

### Latency and feeding time (food provision phase only)

Individuals who approached the food, were considered for the latency comparisons (*N* = 60). We excluded the HIT condition from the analysis as only one dog approached and obtained the food reward. Individuals showed different latencies in the three conditions while approaching for the food (Kruskal – Wallis test, χ^2^ = 34.011, df = 2, p < 0.0001, see Supplementary **Figure S3**). In the FC condition, the dogs approached faster than the NC (Mann – Whitney U test, U = 452.000, df1 = 17, df2 = 30, p < 0.0001) and LIT (Mann – Whitney U test, U = 374.000, df1 = 30, df2 = 13, p < 0.0001) conditions. Latencies were comparable in the NC and LIT conditions (Mann – Whitney U test, U = 156.500, df1 = 17, df2 = 13, p = 0.053).

We found one individual in the FC condition that approached but did not obtain the reward. Thus, we removed the data point for the analysis of feeding time (*N* = 59). We found a significant difference in feeding time (Kruskal – Wallis test, χ^2^ = 8.366, df = 2, p = 0.015, see Supplementary **Figure S4**). Post-hoc pairwise comparisons further revealed a significant difference between feeding times of FC and LIT conditions (Mann – Whitney U test, U = 298.000, df1 = 29, df2 = 13, p = 0.002). Short feeding time in the LIT condition (2.77 ± 0.72 sec) compared to FC (4.58 ± 2.02 sec) might be an indication of dogs’ insecurities due to negative human influence, leading to faster consumption of the food reward. Moreover, we did not see any difference between the other two comparisons (see Supplementary **Table S1**).

## DISCUSSION

Our results underline the free-ranging dogs’ behavioral plasticity in the context of interactions with unfamiliar humans. Dogs adjusted their behavior and showed situation-relevant response to the social cues. Overall, they exhibited a tendency to approach more when food was provisioned compared to the social cue phases, emphasizing the dependence on humans for sustenance. However, comparable but higher levels of approach in the FC condition identified an important role of positive social actions from humans, in order to encourage the initiation of an affiliative relationship. The comparatively higher duration of human proximity in the FC further strengthens this statement. The influence of low-impact threatening cues was very momentary while the effect of the high-impact threatening cues remained even when food was provided. Moreover, dogs avoided the unfamiliar human (E1) with a comparatively higher distance in HIT compared to LIT. Thus, dogs were able to distinguish between the impacts of the threatening cues and responded accordingly, illustrating an optimized strategy. Additionally, the initial reactions and demeanors were consistent with the differential approach of dogs to the corresponding cues.

Apart from showing affiliative responses in the FC condition, dogs showed adjustments and plasticity in their anxious or fearful responses during the threatening cue conditions. Gazing behavior in SCP was predicted by LIT and HIT conditions as dogs gazed more, probably indicating their hesitant nature to approach and also gauging human intentions. On the other hand, FC, LIT and HIT conditions predicted the gazing response in FPP. It is important to note that the short duration of gazing at E1 in the FC condition could be the linked to dogs’ certainty due to affiliative human action. This was also supported by a significantly faster approach to the food. Moreover, dogs depicted a tendency to spend more time while feeding in the FC condition compared to LIT. Thus, the free-ranging dogs acted very specifically in the different conditions displaying a range of social responses that had a high degree of parity with the social cue provided in the experiment.

Free-ranging dogs live in human dominated environments and heavily depend on humans for food (Bhadra et al., 2016; Bhadra and Bhadra, 2014). Apart from scavenging, they directly beg for food from humans (Bhadra and Bhadra, 2014; Sen Majumder et al., 2014). However, getting or retrieving food items can lead to consequences like beating and harassment, which probably have made these dogs opportunistic. In order to avoid negative human impact and maximize the success of getting food, dogs need to identify reliable humans. General avoidance of direct physical contact with unfamiliar humans (Bhattacharjee et al., 2017b) may be a process involved in the same strategy. However, the flexibility might have been achieved by dogs upon receiving positive human reinforcements. For example, adult free-ranging dogs have earlier been shown to adjust their point-following behavior based on reliability of unfamiliar humans (Bhattacharjee et al., 2017a). Interestingly, pet dogs have been shown to trusting unfamiliar humans in a range of scenarios (see review Hare and Woods, 2013), which could possibly be result of solely positive interspecific interactions. Nevertheless, pets are sensitive to behaviors of strangers (Vas et al., 2005), consistent with the results of this study.

Dogs’ differential and situation-specific approach behaviors can be explained by early social interactions with humans (Fox and Stelzner, 1966). The role of domestication is also undeniable, which facilitated dogs’ understanding and sensitivity towards human social cues. It has been shown that even hand-reared wolves (*Canis lupus lupus*), being the closest ancestors of modern day dogs failed to adjust behaviors while interacting with humans in ambiguous situations (Bradshaw and Nott, 1995). Thus, an interplay of domestication and factors like living environment and experience with humans might be the basis of the overall outcomes.

The current experimental design only evaluates dogs’ understanding and sensitivity of human social actions. One potential short-coming of the study was not being able to track individuals and failing to incorporate factors like frequency of positive and negative interactions with humans in their daily lives. Follow-up studies in different geographic regions with varying levels of human influences could be done to see the larger picture. On the brighter side, this study has helped to identify a key element in the ecology of the dog-human relationship, the ability of the dogs to assess a social cue (and thus intent) of unfamiliar humans, which explains why dogs are one of the most successful species in sharing the same niche with humans.

## AUTHOR CONTRIBUTIONS

DB and AB conceived and designed the study. DB and SS performed the field experiments as E1 and E2 respectively. DB coded the videos and analysed the data. The initial draft was written by DB and later reviewed and edited by AB. AB supervised the work. All authors gave final approval for publication.

## ACKNOWLEDGEMENTS

The authors would like to thank Indian Institute of Science Education and Research – Kolkata (IISER – K) for providing infrastructural support.

